# Repair oligodendrocytes demyelinating and disintegrating damaged axons after injury

**DOI:** 10.1101/2023.05.18.541273

**Authors:** Gianluigi Nocera, Adrien Vaquié, Nadège Hertzog, Katharina Steil, Santiago Luis Cañón Duque, Johannes Miedema, Cansu Bagin, Margaryta Tevosian, Beat Lutz, Azadeh Sharifi-Aghili, Katharina Hegner, Doris Vollmer, Seokyoung Bang, Seung-Ryeol Lee, Noo Li Jeon, Stephen M Keyse, Sofía Raigón López, Claire Jacob

**Affiliations:** Institute of Developmental Biology and Neurobiology, Faculty of Biology, Johannes Gutenberg University Mainz, Mainz, Germany; Faculty of Biology, University of Fribourg, Fribourg, Switzerland; Institute of Physiological Chemistry, University Medical Center of the Johannes Gutenberg University Mainz, Mainz, Germany; Max Planck Institute for Polymer Research, Mainz, Germany; School of Mechanical and Aerospace Engineering, Seoul National University, Seoul, South Korea; Jacqui Wood Cancer Centre, Division of Cellular and Systems Medicine, School of Medicine, University of Dundee, Dundee, United Kingdom; Leibniz Institute for Resilience Research (LIR), Mainz, Germany; Department of Biomedical Engineering, Dongguk University, Goyang 10326, Republic of Korea

**Keywords:** Schwann cells, Oligodendrocytes, Dusp6, Axon disintegration, Demyelination, Myelin clearance, Axonal regrowth, Spinal cord injury

## Abstract

After a spinal cord injury, axons fail to regrow, which results in permanent loss of function^1^. This is in contrast with peripheral axons that can regrow efficiently after injury^2^. These differences are partly due to the different plasticity of myelinating cells, Schwann cells and oligodendrocytes, in these two systems^3^. The molecular mechanisms underlying this different plasticity remain however poorly understood. Here, we show that the phosphatase Dusp6^4^ is a master inhibitor of oligodendrocyte plasticity after spinal cord injury. Dusp6 is rapidly downregulated in Schwann cells and upregulated in oligodendrocytes after axon injury. Simultaneously, the MAP kinases ERK1/2 are activated and the transcription factor c-Jun is upregulated in Schwann cells^5,6^, but not in oligodendrocytes. Ablation or inactivation of Dusp6 induces rapid ERK1/2 phosphorylation, c-Jun upregulation and filopodia formation in oligodendrocytes, leading to mechanically-induced, fast disintegration of distal ends of injured axons, myelin clearance and axonal regrowth. Together, our findings provide understanding of the mechanisms underlying the different plasticity of Schwann cells and oligodendrocytes after injury and a method to convert mature oligodendrocytes exhibiting inhibitory cues for axonal regrowth into repair oligodendrocytes reminiscent of repair Schwann cells. We show that repair oligodendrocytes successfully increase the compatibility of the spinal cord environment with axonal regrowth after injury, suggesting a potential use of repair oligodendrocytes as future therapeutic approach to treat spinal cord injuries.

## Main

After a spinal cord injury, axonal regrowth is highly inefficient. This often leads to permanent loss of function that can greatly alter the quality of life of affected individuals. In contrast, traumatic injuries of the peripheral nervous system (PNS) can be efficiently repaired. The reasons for these different regenerative properties are multifold: 1/ PNS axons have in general a higher intrinsic capacity for regrowth as compared to central nervous system (CNS) axons^7^, 2/ in the CNS, a glial scar rapidly forms in the lesion site and acts as a barrier for axonal regrowth^8^, 3/ the myelin of oligodendrocytes (OLs) in the CNS contains several growth-inhibitory factors for axons and thus inhibits axonal regrowth^3^, 4/ axonal fragments in the CNS persist a long time after injury and also act as inhibitors of axonal regrowth^9-11^, 5/ in contrast to the PNS, there is no known guidance mechanism after lesion in the CNS allowing axons to reconnect to their former target^12^, 6/ remyelination in the CNS, which is mostly achieved by OL precursor cells that need to migrate to the lesion site and be in sufficient number to remyelinate axons^13^, may be more challenging than remyelination in the PNS, which is achieved by Schwann cells (SCs) that are already in contact with or are present in the vicinity of damaged axons^2^. In light of the multiple barriers to functional regeneration in the CNS, the combination of several time-controlled regenerative strategies appears to be the most promising approach. Here, we have focused on removing axonal regrowth inhibitory cues that are due to the persistence of axonal and myelin debris distal to the lesion site.

In the PNS, distal ends of injured axons that are disconnected from the neuronal cell body (distal cut axons) disintegrate rapidly after injury^14-17^. We have shown that SCs play an essential role in this process by forming constricting actin spheres around distal cut axons to accelerate their disintegration^17^. In addition, myelinating SCs actively demyelinate by myelinophagy, an autophagic process specific to myelin that is driven by the transcription factor c-Jun^18^. Extracellular signal-regulated kinase (ERK) signaling is also importantly involved in this process^19^. Rapid axon disintegration and demyelination are key events to ensure fast clearing of axon and myelin fragments and removal of growth-inhibitory cues from the injured nerve, and thereby to create a favorable environment for axonal regeneration. In the spinal cord, axonal disintegration after injury is a lot slower than in the PNS^9,10,14,17^. The persistence of axon fragments has been shown to delay axonal regrowth in the PNS and decrease axonal sprouting in the CNS^20-22,11^. Therefore, accelerating the disintegration of distal cut axons has the potential to facilitate axonal regrowth after injury. In contrast to SCs, OLs in contact with damaged axons fail to demyelinate and their myelin that contains several growth-inhibitory factors for axons, prevents axonal regrowth^3^. Promoting the removal of myelin debris and inducing demyelination of OLs in contact with damaged axons distal to the lesion site has thus also the potential to enhance axonal regrowth in the CNS.

In this study, we have identified the phosphatase Dusp6 as a major negative regulator of OL plasticity after axonal lesion. The main known function of Dusp6 is to dephosphorylate and thereby inactivate ERK1/2^4^. In addition, Dusp6 has been shown in some instances to dephosphorylate other substrates including c-Jun-N-terminal kinases (JNKs)^23-25^. Here, we show that the ablation of Dusp6 in mature OLs enables ERK1/2 phosphorylation and c-Jun upregulation upon axonal lesion. This induces a pro-regenerative behavior in OLs similar to repair SCs after a PNS lesion, leading to fast disintegration and demyelination of distal cut axons, and axonal regeneration.

### Injury-induced opposite regulations in Schwann cells and oligodendrocytes

We previously set up and validated microfluidic lesion models of myelinated systems^17^. With these models, we analyzed by RNA sequencing mRNA regulations in SCs at 1 day post axonal lesion (dpl) compared to unlesioned cultures^17^. Among the regulated genes revealed by this analysis, we decided to investigate the potential involvement of the phosphatase Dusp6 that we found downregulated in SCs in our microfluidic lesion models^17^ and also at 1 day post sciatic nerve crush lesion (SNCL) (Fig. 1a), whereas Dusp6 was upregulated in mature OLs at 1 day post spinal cord hemisection injury (SCI) (Fig. 1b) in mice. Dusp6 upregulation in OLs persisted for 5 days after SCI and levels were decreased at 7 days post SCI (Supplementary Fig. 1a). ERK1/2 phosphorylation was rapidly increased after SNCL, already at 1 dpl (Supplementary Fig. 1b), but started only at 7 dpl after SCI (Fig. 1c). ERK1/2 activation is necessary to induce SC demyelination early after peripheral nerve injury but sustained activation of ERK1/2 signaling prevents SC re-differentiation into myelinating SCs and thereby impairs remyelination of regenerated axons^19^. We show here that after a SNCL, ERK1/2 phosphorylation peaks at 3 dpl and then decreases to nearly normalized levels (similar to uninjured nerves) by 12 dpl (Supplementary Fig. 1b). Consistent with a transient activation of ERK1/2 signaling, Dusp6 levels, after initial downregulation at 1dpl (Fig. 1a), increased again with a peak at 3 dpl and nearly normalized levels by 5 dpl (Supplementary Fig. 1c). We and others have previously shown that the transcription factor c-Jun, which drives SC demyelination together with ERK1/2 signaling and the conversion of mature SCs into repair SCs upon axonal lesion^6^, is rapidly upregulated in SCs after peripheral nerve injury^26,27^. Simultaneously, JNKs and c-Jun are phosphorylated^26^. In contrast, we found that c-Jun was downregulated and the levels of phosphorylated c-Jun decreased in mature OLs after a SCI (Fig. 1d). Taken together, these data show that SCs and OLs react radically differently to injury, with SCs rapidly activating a repair program correlating with Dusp6 downregulation, whereas OLs do not activate this repair program and instead rapidly upregulate Dusp6 expression.

**Fig. 1:**
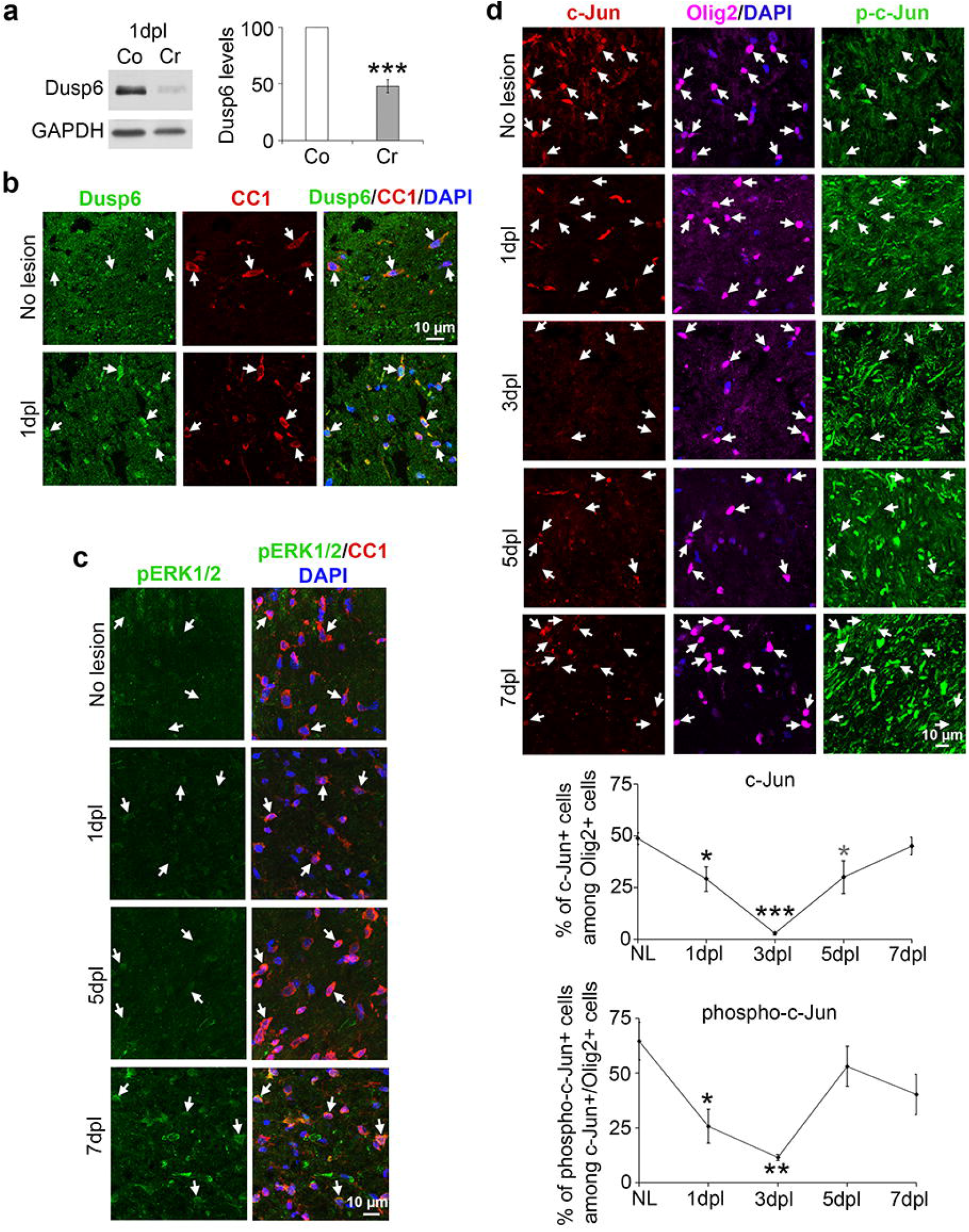
The Dusp6/ERK/c-Jun axis is oppositely regulated in SCs and OLs after injury. **a**, Dusp6 Western blot and quantification normalized to GAPDH showing downregulation of Dusp6 at 1dpl in crushed (Cr) as compared to contralateral (Co) sciatic nerves of adult mice. Paired two-tailed Student’s t-tests, p value: ***<0.001, values=mean, error bars=s.e.m., n=3 animals per group. **b**, Confocal images (z-series projections) of Dusp6 and CC1 (mature OL marker) co-immunofluorescence and DAPI (nuclei) labeling at 1 day post lesion (dpl) and in unlesioned (No lesion) mouse spinal cords. **c**, Confocal images (z-series projections) of phospho-ERK1/2 (pERK1/2) and CC1 co-immunofluorescence and DAPI labeling at 1, 5 and 7 dpl and in unlesioned mouse spinal cords. **d**, Confocal images (z-series projections) of c-Jun, phospho-c-Jun (p-c-Jun) and Olig2 (OL marker) co-immunofluorescence and DAPI labeling at 1, 3, 5 and 7 dpl and in unlesioned mouse spinal cords, and percentage of c-Jun-positive cells among Olig2-positive cells and of phospho-c-Jun-positive cells among c-Jun/Olig2-double positive cells. Unpaired one-tailed (grey asterisk) or two-tailed (black asterisks) Student’s t-tests, p value: *<0.05, **<0.01,***<0.001, values=mean, error bars=s.e.m., n=3 animals per time point (73 to 181 Olig2-positive cells counted per animal). Arrows show CC1 or Olig2-positive cells (**b**,**c**,**d**).

### Axon disintegration and regrowth in myelinated cultures after lesion

To investigate the function of Dusp6 in OLs after injury, we first used our microfluidic lesion models of neuron/OL co-cultures^17^. At the myelinated culture stage, we downregulated Dusp6 specifically in OLs by using a lentiviral vector carrying a highly efficient Dusp6-specific shRNA or a non-targeting control shRNA (Supplementary Fig. 2a). We previously showed in this microfluidic system that virtually all OLs are efficiently transduced by using highly concentrated lentiviruses in the OL compartment, while neurons do not get transduced or rarely (0-2 neurons per device)^17,28^. After laser axotomy, the majority of regrowing axons rapidly collapsed and axonal regrowth was overall of low efficiency and over short distance in cultures where OLs received the control shRNA. In comparison, in cultures where OLs received the Dusp6 shRNA, less axons collapsed and axons regrew overall more efficiently, some over long distance (>500 μm) after lesion (Fig. 2a). We showed that Dusp6 is rapidly downregulated in SCs after lesion (Ref. 17 and Fig. 1a). We then asked whether Dusp6 downregulation is involved in SC pro-regenerative behavior after lesion. To answer this question, we used our microfluidic lesion models of neuron/SC co-cultures and we transduced SCs with a lentiviral vector expressing Dusp6 (Supplementary Fig. 2b) to prevent Dusp6 downregulation after laser axotomy. We found that axonal regrowth was significantly slower in cultures where SCs were transduced with the Dusp6-expressing lentivirus as compared to cultures where SCs received a control lentivirus (Fig. 2b). In addition, the disintegration of distal cut axons was accelerated in cultures where Dusp6 was downregulated by shRNA in OLs compared to cultures where OLs were transduced with a lentivirus carrying a control shRNA (Fig. 2c). These data indicate that Dusp6 downregulation indeed contributes to SC pro-regenerative behavior and in OLs can lead to rapid disintegration of distal cut axons and prevent OL-mediated inhibition of axonal regrowth after lesion.

**Fig. 2:**
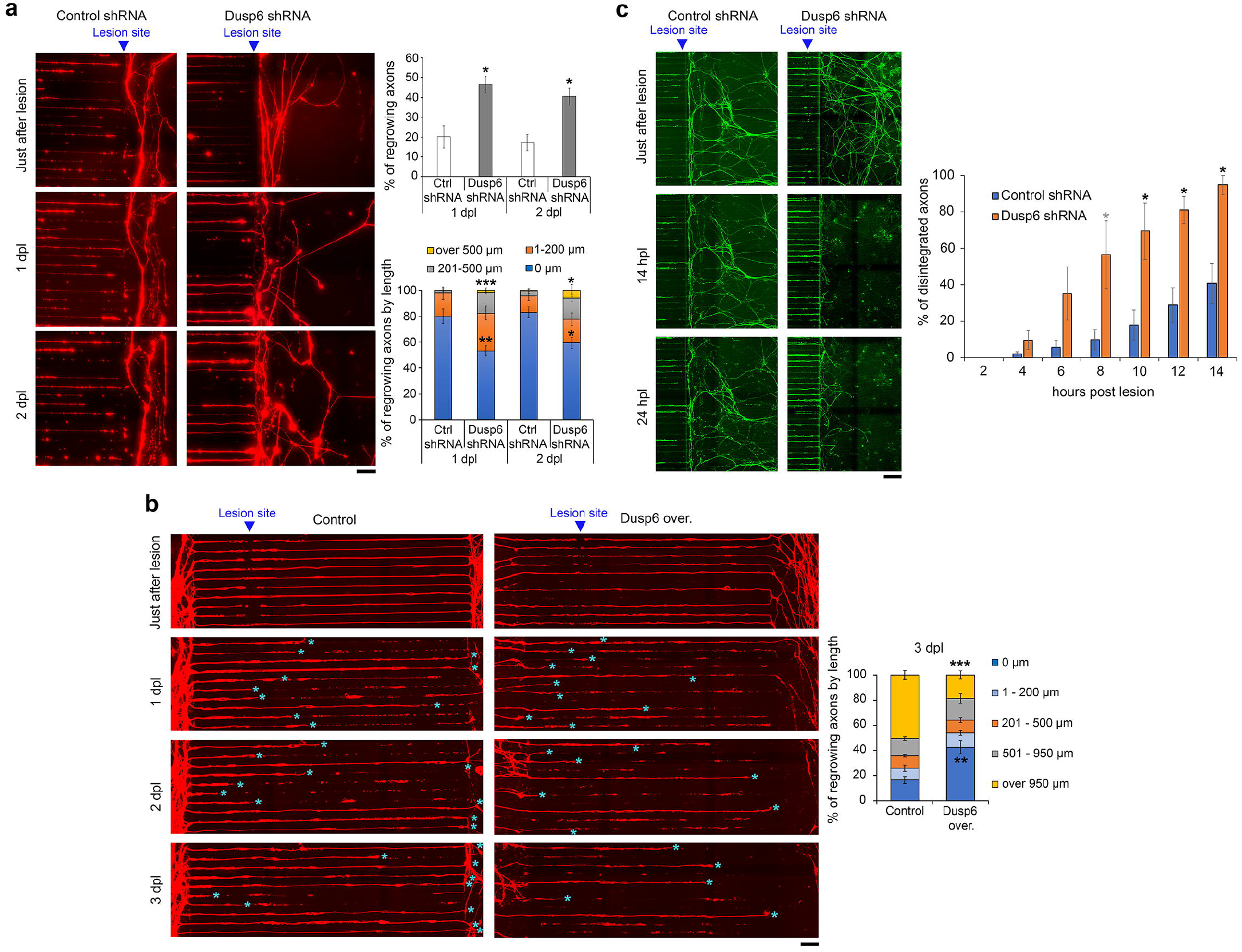
Dusp6 prevents distal cut axon disintegration and axonal regrowth. **a**,**c**, Time-lapse imaging (wide-field) at different time points after lesion of DsRed-labeled (**a**) or GFP-labeled (**c**) axons in microgrooves of neuron/OL cultures where OLs were transduced with lentiviruses carrying either a Dusp6-specific shRNA or a non-targeting control shRNA. Note that lesions were carried out at the exit of microgrooves to quantify axonal regrowth (**a**) and the percentage of disintegrated distal cut axons (**c**) in the chamber where OLs interact with axons. In (**a**), the upper graph shows the percentage of regrowing axons and the lower graph shows the length distribution of axonal regrowth after lesion. **b**, Time-lapse imaging (wide-field) at different time points after lesion of DsRed-labeled axons in microgrooves of neuron/SC cultures where SCs were transduced with lentiviruses expressing Dusp6 (Dusp6 over.) or with control lentiviruses (empty vector), and graph showing the length distribution of axonal regrowth after lesion. Blue asterisks indicate the position of the tip of axons in the microgrooves and axons that have regrown beyond at their exit of the microgrooves. (**a**,**b**,**c**) Fifty-two to 68 axons (**a**), 76 to 192 axons (**b**), and 27 to 81 axons (**c**) were quantified per chamber per day, n=4-6 chambers per group, unpaired one-tailed (grey asterisk) or two-tailed (black asterisks) Student’s t-test, p value: *<0.05, **<0.01, ***<0.001, values=mean, error bars=s.e.m. Scale bar, 50 μm (**a**,**b**), 100 μm (**c**).

At the molecular level, Dusp6 downregulation by shRNA in purified and differentiated primary OLs resulted in increased levels of phosphorylated ERK1/2, as expected (Fig. 3a). To test a potential involvement of increased ERK1/2 activation in the rapid disintegration of distal cut axons and the induction of axonal regrowth resulting from Dusp6 downregulation in OLs, we used the specific ERK1/2 inhibitor MK-8353. We found that ERK1/2 inhibition in the OL chamber slowed down distal cut axon disintegration and impaired axonal regrowth induced by Dusp6 downregulation in OLs (Fig. 3b). We have previously shown that SCs react to axonal lesion by building constricting actin spheres along distal cut axons to accelerate their disintegration, which leads to fast axon debris clearance and fast axonal regrowth^17^. ERK1/2 activation is known to promote actin dynamics^29^. We thus hypothesized that Dusp6 downregulation leads to accelerated distal cut axon disintegration through increased actin polymerization around distal cut axons. In our microfluidic models, virtually all OLs can be transduced with 2 different high-titer lentiviruses at a 2-day interval between the 2 lentiviruses^17^. We thus first transduced OLs with lentiviral vectors expressing the Dusp6 shRNA and 2 days later with lentiviral vectors expressing Lifeact-GFP, which dynamically labels F-actin^17^. Interestingly, the shape of OLs where Dusp6 was downregulated by shRNA was altered and actin polymerisation rapidly increased after axotomy, whereas the shape of OLs transduced with the control shRNA lentivirus remained similar and there was no detectable actin polymerisation changes (Fig. 3c). Remarkably, while OLs transduced with the control shRNA lentivirus remained static after lesion with only occasional minor actin process movements and did not form filopodia, OLs transduced with the Dusp6 shRNA lentivirus displayed major actin process changes after lesion, from a flat shape with a few filopodia to a contracted spherical actin shape exhibiting many filopodia (Fig. 3c). Filopodia are actin structures that sense the environment and have the capacity to make adhesive contact with the extracellular matrix, pathogens and adjacent cells, and subsequently exert pulling forces^30^. Accordingly, time-lapse imaging shows that OLs transduced with the Dusp6 shRNA lentivirus pull distal cut axons until their disintegration (Fig. 3c and Supplementary Video 1). After disintegration of distal cut axons, actin filaments in OLs transduced with the Dusp6 shRNA lentivirus depolymerized completely (Fig. 3c).

**Fig. 3:**
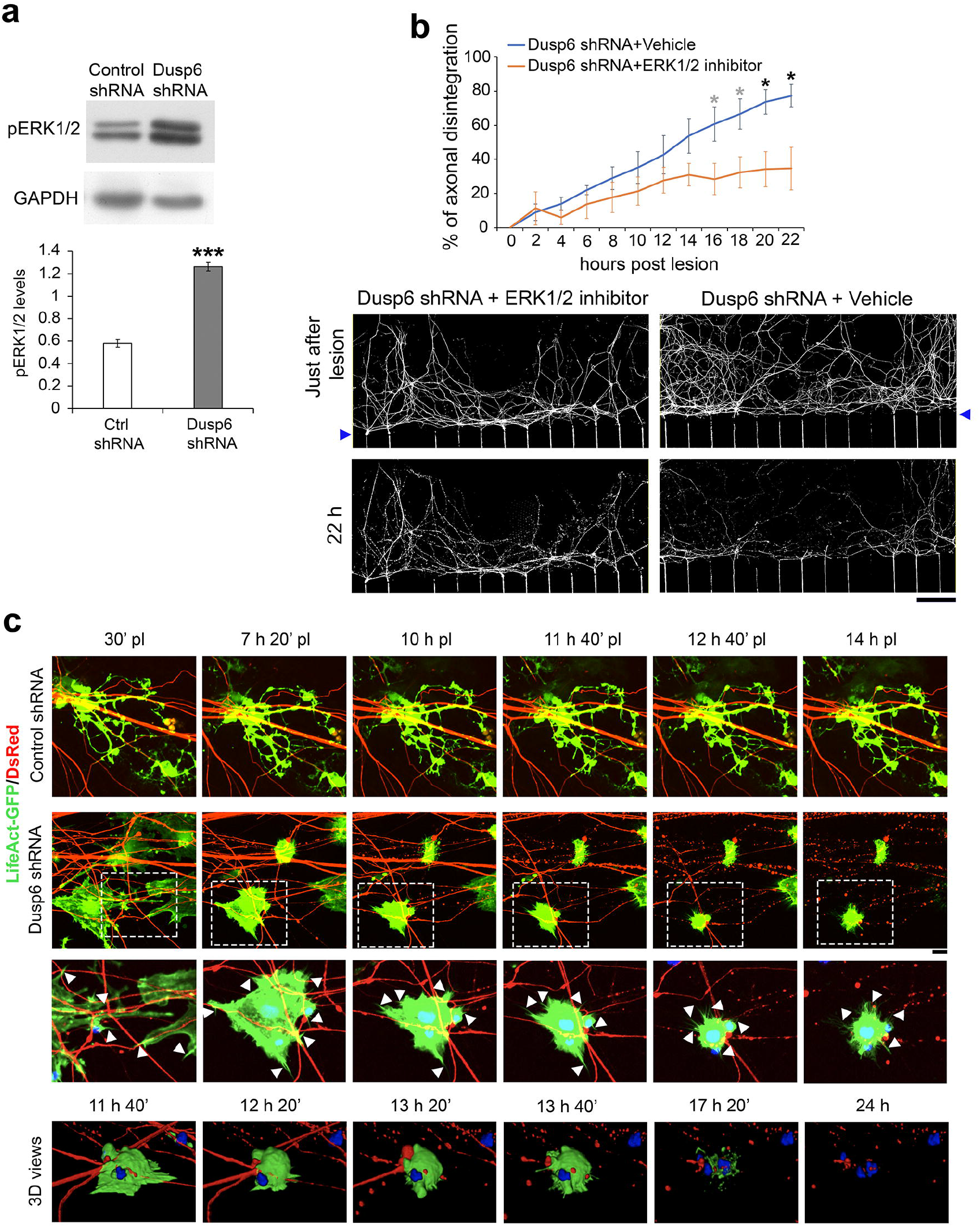
Dusp6 downregulation in OLs induces ERK-dependent distal cut axon disintegration and actin dynamics. **a**, phospho-ERK1/2 (pERK1/2) Western blot and quantification normalized to GAPDH showing increased pERK1/2 levels in primary differentiated OLs transduced with a Dusp6-specific shRNA lentivirus compared to a non-targeting control shRNA lentivirus. Unpaired two-tailed Student’s t-tests, p value: ***<0.001, values=mean, error bars=s.e.m., n=3 independent experiments. **b**, Time-lapse imaging (wide-field) at different time points after lesion of DsRed-labeled axons in microgrooves and chamber #2 of neuron/OL cultures where OLs were transduced with lentiviruses carrying a Dusp6-specific shRNA and treated either with the specific ERK1/2 inhibitor MK-8353 or its vehicle, and quantification of distal cut axon disintegration every two hours for 22 hours. The integrated density (mean grey value) of labeled axons in the 3 same ROI per chamber was averaged at each time point, n=3 chambers per group, unpaired one-tailed (grey asterisks) or two-tailed (black asterisks) Student’s t-test, p value: *<0.05, values=mean, error bars=s.e.m. Scale bar, 100 μm. **c**, Time-lapse confocal imaging at different time points after lesion of DsRed-labeled axons and LifeAct-GFP-labeled OLs (+ NucBlue™ Live ReadyProbes™ Reagent shown in magnifications of Dusp6 shRNA-transduced OLs to label nuclei) in chamber #2 of neuron/OL cultures where OLs were transduced with lentiviruses carrying either a Dusp6-specific shRNA or a non-targeting control shRNA. The two upper rows are z-series projections, the third row is a magnification of the region highlighted by a dashed white box in the second row, and the 4^th^ row shows 3D views at different time points starting from 11 h 40’ after lesion of the region highlighted by a dashed white box. White arrowheads point to filopodia structures. Scale bar=10 μm.

ERK1/2 activation has been shown to upregulate c-Jun^19,31^. Remarkably, Dusp6 downregulation by shRNA in purified and differentiated primary OLs resulted in strongly increased c-Jun levels (Fig. 4a). Consistently, Dusp6 overexpression in SCs led to a decreased percentage of c-Jun-positive SCs after axonal lesion (Supplementary Fig. 3). We thus hypothesized that Dusp6 downregulation in OLs leads to increased c-Jun levels due to increased ERK1/2 activation, and thereby induces a SC-like repair phenotype in OLs after lesion. Indeed, in purified and differentiated primary OL cultures, c-Jun upregulation induced by Dusp6 downregulation was prevented by ERK1/2 inhibition, while JNK inhibition had a milder but significant effect on c-Jun levels (Fig. 4a). In addition to increasing phospho-ERK1/2 levels, Dusp6 downregulation also increased phospho-JNK levels and this was prevented by ERK1/2 inhibition (Fig. 4a), indicating that the increase of phospho-JNK levels by Dusp6 downregulation is mediated by ERK1/2 activation. Remarkably, Dusp6 downregulation led to a strong downregulation of MBP expression, which was prevented by ERK1/2 inhibition and to lesser extend by JNK inhibition (Fig. 4a). To test whether Dusp6 downregulation in OLs leads to accelerated distal cut axon disintegration and to the induction of axonal regrowth through increased c-Jun levels, we first transduced OLs with lentiviral vectors expressing Dusp6 shRNA and second with lentiviral vectors expressing a c-Jun-specific shRNA or a non-targeting control shRNA. We show here that c-Jun downregulation in OLs slows down axonal regrowth but does not affect distal cut axon disintegration induced by Dusp6 downregulation (Fig. 4b). Taken together, these data show that Dusp6 downregulation allows rapid ERK1/2 activation, leading to c-Jun upregulation and to the acquisition of a SC-like repair phenotype. Interestingly, ERK1/2 activation induced by Dusp6 downregulation promotes axonal regrowth after lesion at least partially via c-Jun upregulation, and induces the disintegration of distal cut axons in a c-Jun-independent manner.

**Fig. 4:**
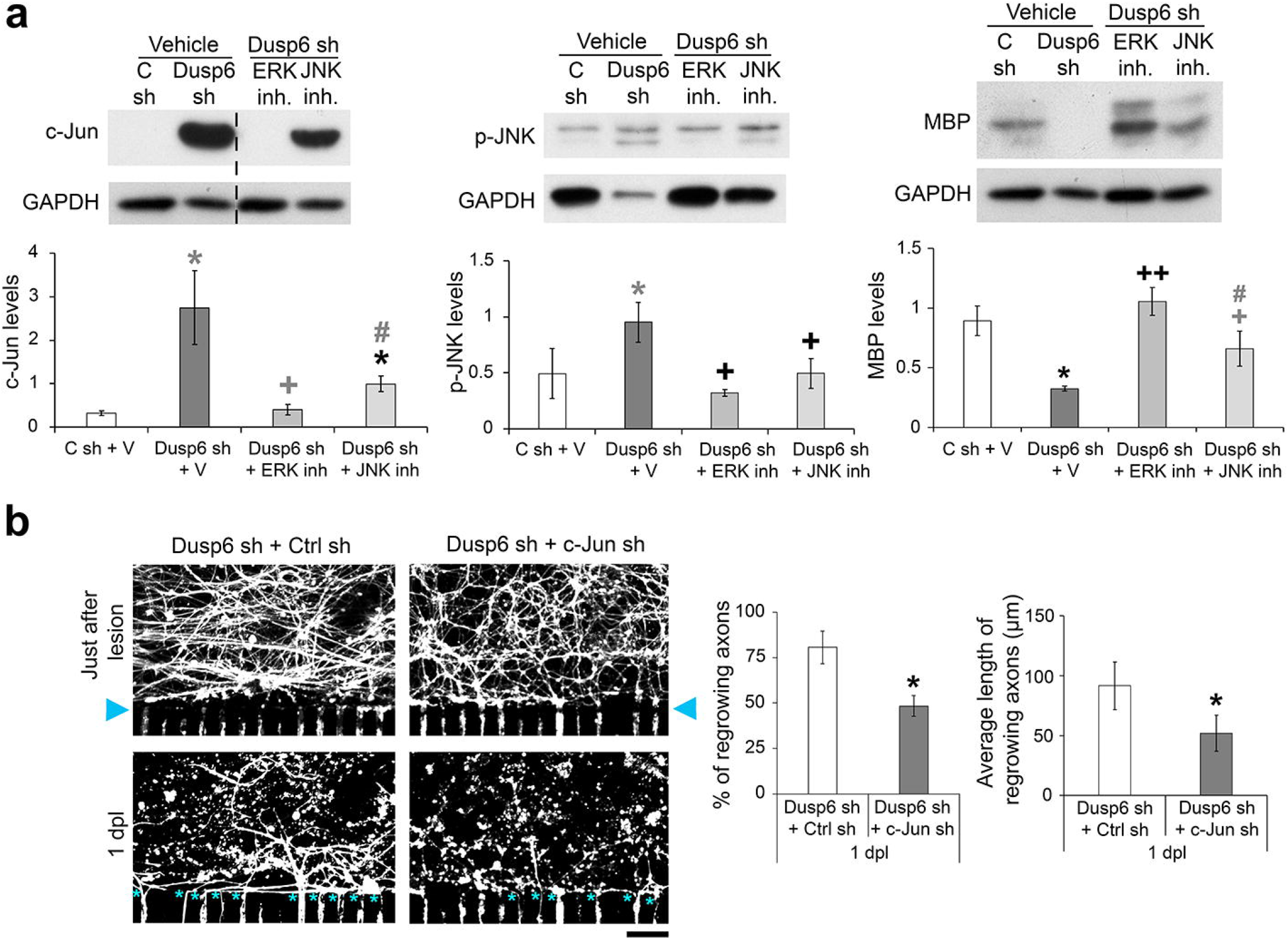
Dusp6 downregulation in OLs induces c-Jun-dependent axonal regrowth. **a**, c-Jun, phospho-JNK (p-JNK) and MBP Western blots and quantification normalized to GAPDH in primary differentiated OLs transduced with a Dusp6-specific shRNA lentivirus (Dusp6 sh) or a non-targeting control shRNA lentivirus (C sh) and treated with the specific ERK1/2 inhibitor MK-8353 (ERK inh) or the specific JNK1/2 inhibitor JNK-IN-8 (JNK inh) or their vehicle (V). Unpaired or paired (pJNK: Dusp6 sh/C sh and Dusp6 sh + JNK inh / Dusp6 sh + V; MBP: Dusp6 sh + JNK inh /Dusp6 sh + ERK inh) one-tailed (grey asterisks, crosses or hashtags) or two-tailed (black asterisks or crosses) Student’s t-tests, p value: *,+,#<0.05, ++<0.01; asterisks indicate significance compared to C sh + V, crosses indicate significance compared to Dusp6 sh + V, hashtags indicate significance compared to Dusp6 sh + ERK inh; values=mean, error bars=s.e.m., n=3 independent experiments. **b**, Wide-field imaging just after lesion and at 1 dpl of DsRed-labeled axons in microgrooves of neuron/OL cultures where OLs were transduced with lentiviruses carrying a Dusp6-specific shRNA and either a non-targeting control shRNA or a c-Jun-specific shRNA, and graphs showing the percentage and the average length of regrowing axons. Lesions were carried out near the exit of the microgrooves (blue arrowheads). Blue asterisks indicate regrowing axons. Fourty-three to one-hundred axons were quantified per chamber per time point, n=3 chambers per group, unpaired (% of regrowing axons) or paired (average length of regrowing axons) two-tailed Student’s t-test, p value: *<0.05, values=mean, error bars=s.e.m. Scale bar, 50 μm.

### Axon disintegration and regrowth after spinal cord injury in mice

Next, we aimed at translating our findings obtained with our microfluidic cell culture models to an *in vivo* model of SCI. We asked whether repair OLs can be induced *in vivo* and if this is the case, whether these repair OLs can promote distal cut axon disintegration and axonal regrowth after SCI. To address these questions, we generated a tamoxifen-inducible Dusp6 knockout (Dusp6 KO) mouse line where *Dusp6* is specifically ablated in the CNS in mature OLs by crossing *Dusp6* floxed mice^32^ with *PlpCreERT2* mice^33^ expressing the tamoxifen-inducible CreERT2 recombinase under control of the *Plp* promoter. In some cases, we also crossed Dusp6 KO mice with the R26-stop-EYFP reporter mouse line^34^ to label recombined mature OLs, or with a Thy1-GFP M mouse line^35^ for sparse labeling of different neuronal subsets. As control mice, we used *PlpCreERT2*-negative littermates that were treated with tamoxifen at the same time as Dusp6 KO mice.

Because Dusp6 expression is below detectable levels in mature OLs of unlesioned spinal cords, we did not determine the onset of protein loss after tamoxifen, but instead fixed an arbitrary time point of 2 weeks after tamoxifen injection to carry out a SCI in Dusp6 KO and control mice. We showed above that in wild type mice, Dusp6 is upregulated in mature OLs between 1 and 5 days post SCI (Fig. 1b and Supplementary Fig. 1a) and that phospho-ERK1/2 is not detected in these cells at these time points (Fig. 1c). In Dusp6 KO mice, Dusp6 was efficiently ablated in OLs with no detectable expression at 1 or 5 days post SCI (Fig. 5a). Consistently, strong phosphorylated ERK1/2 levels were detected at 1 day post SCI and were sustained until at least 5 days post SCI in mature OLs of Dusp6 KO mice, while ERK1/2 phosphorylation was not detected at these time points in mature OLs of control mice (Fig. 5a). Similarly, c-Jun-expression was also increased in mature OLs at these time points in Dusp6 KO mice compared to control mice (Fig. 5b). Remarkably, the disintegration of distal cut axons below the lesion site was significantly enhanced already at 3 days post SCI in Dusp6 KO mice compared to control mice (Fig. 5c).

**Fig. 5:**
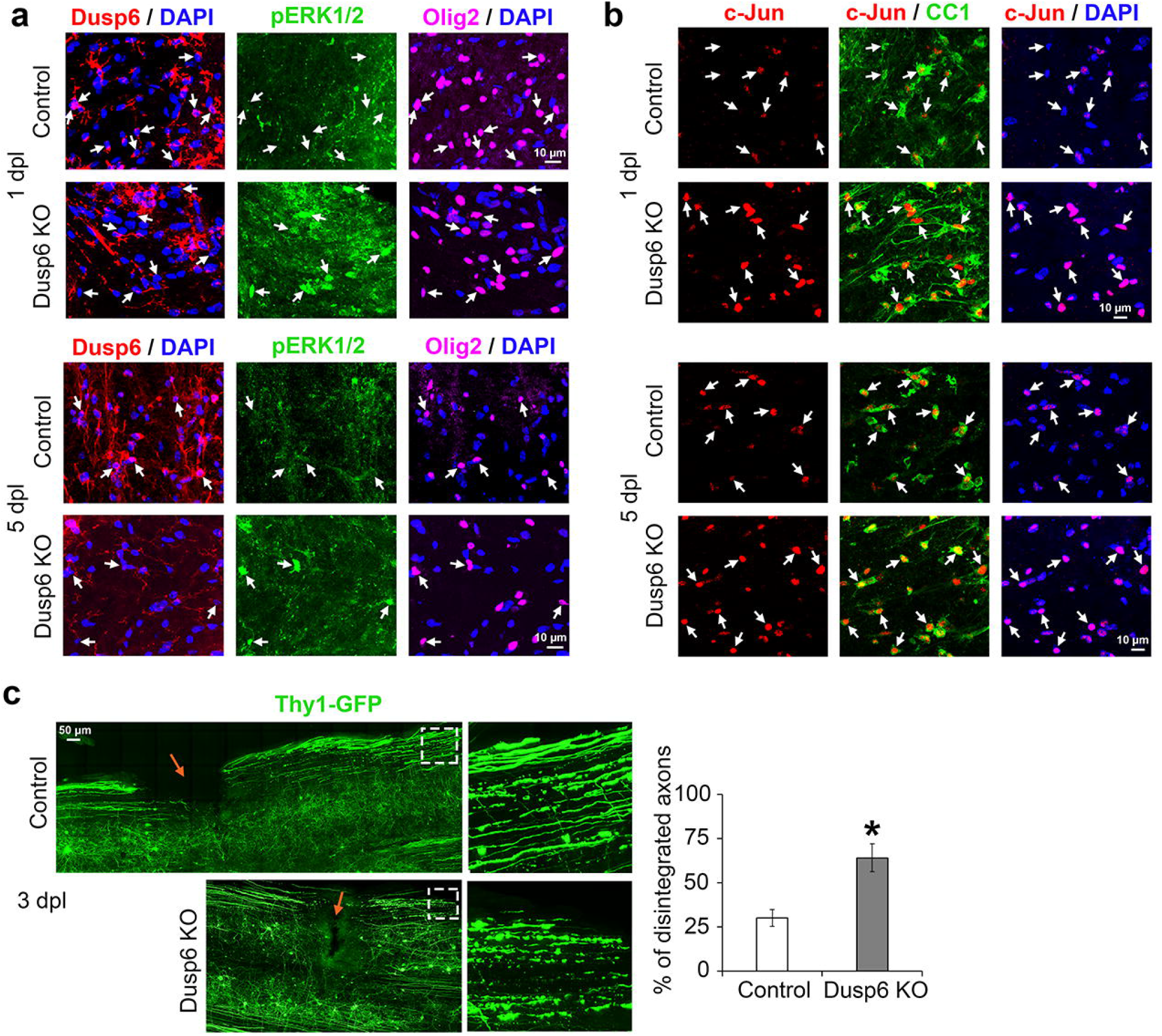
Dusp6 ablation in OLs leads to increased phospho-ERK1/2 and c-Jun levels and to fast distal cut axon disintegration after SCI. **a**,**b**, Confocal images (z-series projections) of phospho-ERK1/2 (pERK1/2), Dusp6 and Olig2 (OL marker) co-immunofluorescence (**a**) or c-Jun and CC1 (mature OL marker) co-immunofluorescence (**b**) and DAPI labeling at 1 and 5 dpl in spinal cords of Dusp6 KO and control mice. White arrows point to OLs. **c**, confocal imaging and z-series projections of Thy1-GFP-labeled neurons at 3 dpl in Dusp6 KO and control mice and quantification of axonal disintegration below the lesion (17-72 counted axons per ROI, 3 ROI per animal quantified below lesion site, n=3 animals per group), unpaired two-tailed Student’s t-test, p value: *<0.05. Orange arrows indicate the lesion site. The spinal cord regions on the right side of the orange arrows are below the lesion site. Images on the right side are magnifications of the regions highlighted by a dashed white box on the left images.

Since c-Jun was upregulated in mature OLs in Dusp6 KO mice as described above, and c-Jun has been shown to induce demyelination in SCs after lesion^18^, we asked whether myelin is also cleared distal to the SCI site in the absence of Dusp6. Indeed, in Dusp6 KO mice, myelin clearance was increased distal to the lesion site compared to control littermate mice at 1 month post SCI (Fig. 6a). Remarkably, several axons had regrown across the glial scar at the lesion site and distal to the lesion site at 1 month post SCI in Dusp6 KO mice, whereas no axon had grown across the lesion site in control littermates of Dusp6 KO mice (Fig. 6b). Taken together, these data indicate that Dusp6 ablation in mature OLs after SCI allows their conversion into repair OLs, reminiscent of repair SCs after a peripheral nerve lesion. In turn, repair OLs promote distal cut axon disintegration, demyelination distal to the lesion site, and axonal regrowth across and below the lesion site.

**Fig. 6:**
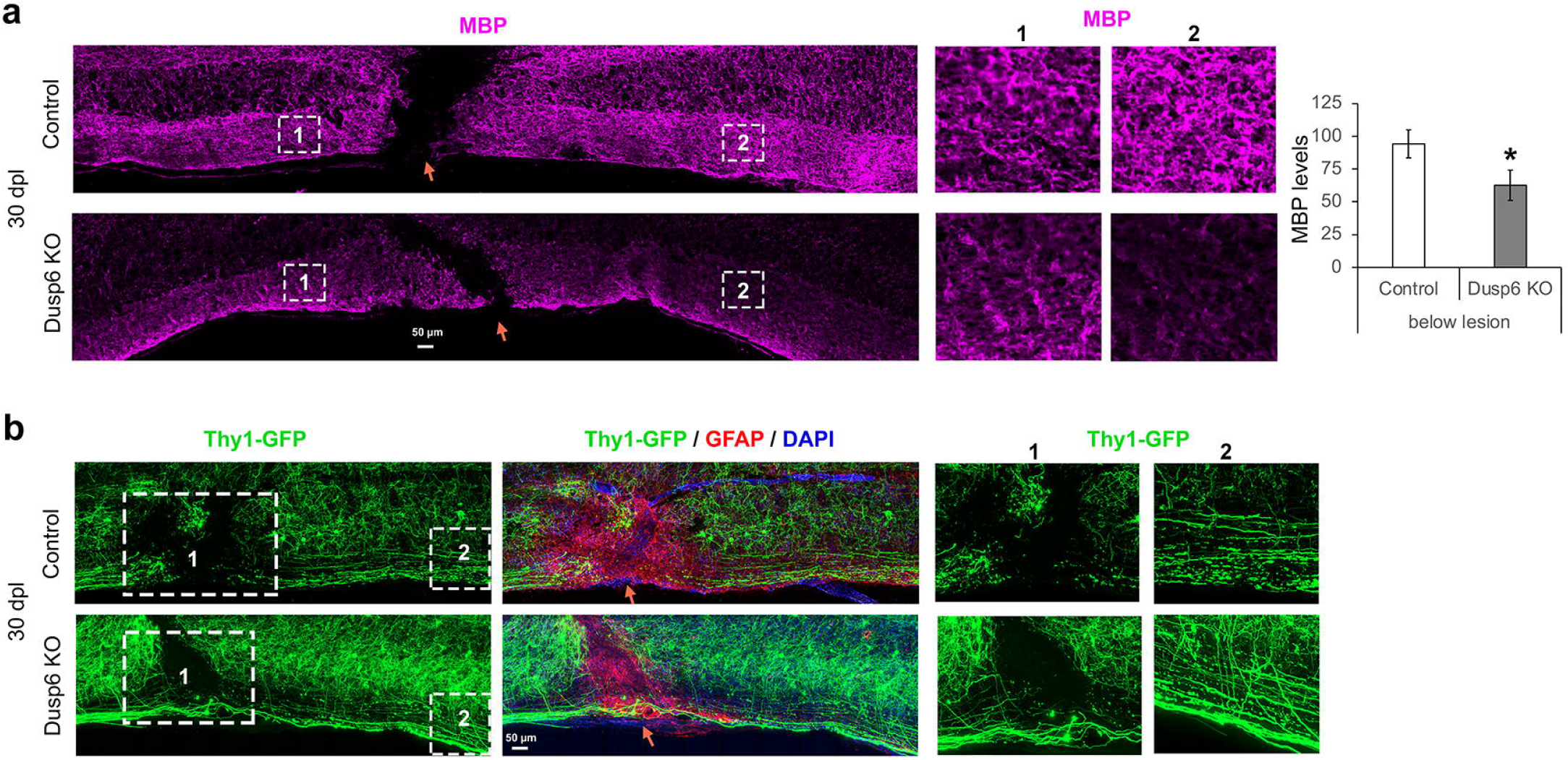
Dusp6 ablation in OLs enables demyelination of damaged axons and axonal regrowth through the glial scar. **a**, Confocal images (z-series projections) of MBP immunofluorescence on Dusp6 KO and control mouse spinal cords at 30 days post SCI (30 dpl) and graph showing the average intensity of MBP signal below the lesion site in the white matter (3 ROI averaged per animal). **b**, Confocal images (z-series projections) of Thy1-GFP-labeled neurons and GFAP (astrocyte/glial scar marker) immunofluorescence and DAPI labeling (nuclei) on Dusp6 KO and control mouse spinal cords at 30 dpl. The GFAP staining delineate the glial scar. Representative images are shown, n=3 animals per group, paired one-tailed Student’s t-test, p value: *<0.05. Images on the right side labeled 1 and 2 are magnifications of the regions highlighted by dashed white boxes on the left images. Orange arrows point to the lesion site. The spinal cord regions on the right side of the orange arrows are below the lesion site.

## Discussion

As explained above, regeneration after SCI requires to overcome multiple barriers^1^. Enormous progress in this domain has however been done in the past 5 years and has shown that biomimetic epidural electric stimulation by electrodes implantation in the spinal cord is an efficient strategy to allow patients with SCI to recover mobility^36-38^. This method is however highly invasive and needs neurostimulation platforms combining ultrafast wireless communication with control units that decode motor intentions^37^. Biology repair strategies constitute alternative approaches to neural engineering and could also be potentially used in combination with neural engineering approaches for synergistic effects^39^.

Injuries of the PNS, in contrast to the CNS, can spontaneously regenerate due to the plasticity of SCs, which convert into repair SCs upon injury to foster axonal regrowth. In contrast, OLs do not convert into repair cells after a CNS injury and instead inhibit axonal regrowth. SCs can enhance the regeneration of both PNS and CNS axons^40^. In addition, transdifferentiation of a fraction of OL precursor cells into PNS-like SCs has been observed after SCI and correlates with partial regeneration and functional recovery^41,42^; this is however limited to lesions of dorsal column axons. This suggests that inducing a SC-like behavior in OLs after injury could promote CNS repair. In this study, we aimed at converting mature OLs into repair OLs after SCI by mimicking regulations occuring in mature SCs leading to their conversion into repair SCs after a PNS injury.

C-Jun and ERK1/2 signaling have been shown to be the main inducers of the SC repair phenotype in the PNS^6,18,19^. We found here that early after lesion, Dusp6 is downregulated in SCs but upregulated in OLs, and that preventing Dusp6 expression in OLs leads to the activation of ERK1/2 and to subsequent upregulation of c-Jun. We show that this is sufficient to induce a SC-like behavior in OLs after lesion. Indeed, ablation of Dusp6 in OLs led to fast disintegration of distal cut axons and enabled axonal regrowth, both in our microfluidic lesion model of neuron/OL cultures and in mice after SCI. In addition, ablation of Dusp6 in OLs resulted in the clearing of myelin debris distal to the SCI site. Thus, inhibitory cues exerted by distal cut axons and OL myelin were removed, creating a favorable CNS environment for axonal regrowth.

Conceptually, this work answers two fundamental questions, the first one being “Can mature OLs convert into repair OLs, reminiscent of repair SCs, upon injury?”, and the second one “Can repair OLs promote axonal regrowth after SCI?”. Here, we positively answer these two questions. In addition, we provide mechanistic understanding underlying the conversion of mature OLs into repair OLs. Future work is needed to develop drugs that downregulate or inhibit Dusp6 specifically in oligodendrocytes below the spinal cord injury site in a timely manner to allow remyelination after axonal regrowth and test whether this strategy promotes functional recovery.

## Methods

### Statistical analyses

For each data set presented, experiments were performed at least 3 times and *p* values were calculated using two-tailed (black asterisks, crosses or hashtags) or one-tailed (grey asterisks, crosses or hashtags) Student’s *t*-tests. *P* values: *<0.05, **<0.01, ***<0.001, values=mean, error bars=s.e.m. Sample size was determined by the minimal number of animals or individual experiment required to obtain statistically significant results and increased in some cases to improve confidence in the results obtained. No animal or data point was excluded from the analysis.

### Animals

To induce ablation of Dusp6 in mature OLs of adult mice, *Dusp6* floxed mice^32^ were crossed with mice expressing a tamoxifen-inducible Cre recombinase under control of the OL-specific *Plp* promoter^33^ (*Plp*CreERT2). To ablate Dusp6, mice received daily injections of 2 mg tamoxifen (Sigma) for five consecutive days. In some cases, these mice were additionally crossed with the R26-stop-EYFP reporter mouse line^34^ to label recombined mature OLs or with a Thy1-GFP M mouse line^35^ to label a fraction of different neuronal subsets. In other cases, the Thy1-GFP M reporter lines was used alone (without other transgene) and in other cases, the reporter line R26-stop-EYFP was crossed to the *Plp*CreERT2 mouse line only. Genotypes were determined by PCR on genomic DNA.

This study complies with all relevant ethical regulations concerning animal use, which was approved by the Veterinary office of the Canton of Fribourg, Switzerland and the Veterinary office (Landesuntersuchungsamt) of Rheinland-Pfalz, Germany.

### Surgical procedures

For all surgical procedures, we used isoflurane (3% for induction, 1.5-2% for narcosis during the operation) for anesthesia. For analgesia, 0.1 mg/kg/body weight buprenorphine (Temgesic; Essex Chemie) was administered by i.p. injection 1 h before surgery and every 4 hours (minimum 2 injections and maximum 3 injections) the day of surgery. The day after surgery 1 ml agarose gel containing 0.027 mg/ml buprenorphine was fed twice a day (morning and evening). Sciatic nerve crush lesions were carried out on 3 to 4-month old adult mice (males and females), as previously described^27^. Spinal cord hemisections were carried out at T8 level on 3 to 4-month old adult mice (males and females) as described^44^. To ensure a complete hemisection, a 36G needle was inserted at the midline of the spinal cord and moved carefully towards the left until the edge of the spinal cord. No randomization method was used, but experimenters were blinded to the experimental group and received only the animal number given at birth by the animal caretaker.

### Microfluidic devices

Our microfluidic lesion models of neuron/SC and neuron/OL cultures were previously described^17,28^.

### DRG neuron/SC myelinated cultures

Dorsal root ganglion (DRG) explant were isolated from embryonic day 14.5 (E14.5) Wistar rat embryos, dissociated and cultured as previously described^17,45^.

### DRG neuron/OL myelinated cultures

DRG explant from E14.5 rat embryos were isolated and neurons were dissociated and cultured in microfluidic devices as previously described^17^.

### Live imaging, laser axotomy and image processing

For live-imaging, neurons were labeled either in red or green by adding to chamber#1 (chamber containing neuronal cell bodies) 0.5-2μL of highly concentrated lentivirus generated as previously described^17^. Red fluorescence was obtained by transducing neurons with lentiviruses expressing DsRed under control of the neuron-specific *Synapsin* promoter. Green fluorescence was obtained by transducing neurons with lentiviruses expressing GFP under control of the CMV promoter. OLs F-Actin structure was labelled in green by adding to chamber#2 0.5-2μL of highly concentrated Lifeact-GFP lentivirus under control of the CMV promoter. Twenty-four hours before carrying out the laser axotomy all medium was replaced with Minimum Essential Medium (MEM). All axons of the device passing through the microgrooves were lesioned with a laser ablation module as described^17^. Laser axotomy was conducted with a 40x or a 60x Water NA1.25 Apochromat objective.

For 2D and 3D time-lapse imaging, we used a VisiScope spinning disk confocal microscope CSU-W1 (Visitron) to acquire wide-field or confocal images at 20, 30 or 60 min intervals on several stage positions (5 to 10 with at least 10% image frame overlap) during 24 to 48 h with a 40x Oil NA1.25 Apochromat objective or at 24 h intervals on 12 to 60 stage positions during 2 to 4 days with a 20x Air NA0.75 PlanApochromat objective. For 3D time-lapse imaging, optical sections of 0.3 to 0.6 μm thickness (between 30 and 100 stacks) were acquired. For confocal imaging, optical sections of 0.6 to 1.5 μm thickness (between 25 and 100 stacks) were acquired. Multiple stage positions were automatically stitched by processing with the Visiview software (Visitron) and different channels were merge with Fiji and/or Adobe Photoshop (CC 20.0.8 Release). Specific Macro were written to convert the saved .stk or .ome.tf2 raw data to RGB mode, create hyperstacks and adjust brightness and contrast using defined minimal and maximal values and convert to .tif files.

### Primary rat oligodendrocyte cultures

Rat primary oligodendrocytes precursors cells (OPCs) were isolated from neonatal rat neocortices and cultured as described^46^. OPCs were first differentiated for 12 days, then the Dusp6 shRNA or non-targeting control shRNA lentiviruses were added and cells were kept in the differentiation medium for 5 additional days before analysis. In some cases, cells were treated for 2 more days in differentiation medium with 1 μM JNK1/2 inhibitor (JNK-IN-8, MCE, #1410880-22-6), 300 nM ERK1/2 inhibitor (MK-8353, MCE, #HY-111407), and then analyzed.

### Primary rat Schwann cell cultures

Primary rat SC cultures derived from P2 Wistar rat sciatic nerves were purified, dissociated and cultured as previously described^27,47^.

### Generation of Lentivirus

Highly concentrated lentiviral particles were produced as previously described (Brügger et al., 2015, Vaquié et al., 2019). Constructs used to produce lentiviruses: packaging constructs pLP1, pLP2 and pLP/VSVG (Invitrogen), pLV-LSyn-RFP^48^ (Addgene construct #22909), pLentiLox 3.7 (ATCC), Lifeact-GFP (kind gift from Dr. Olivier Pertz, University of Bern, Switzerland), Dusp6^49^ (Addgene construct #27975), Dusp6 shRNA (Sigma, mission shRNA, TRCN0000317759: GTTTGGCATCAAGTACATCTT), c-Jun shRNA (Sigma, mission shRNA, TRCN0000229527: GCTAACGCAGCAGTTGCAAAC) or a non-targeting control shRNA (Sigma, SHC001, MISSION pLKO.1-puro Empty Vector Control).

### Immunofluorescence

For immunofluorescence in microfluidic chambers, cells were fixed 30 min with 4 % paraformaldehyde (PFA, Sigma) at room temperature (RT), and washed 3 times 15 min with PBS. Cells were subsequently blocked for 3 hrs with blocking buffer (0.3 % Triton X-100, 5 % BSA, PBS), and then incubated for 2 days at 4°C with primary antibodies in blocking buffer. Cells were then washed three times 30 min with blocking buffer and incubated for 6 hrs in the dark with secondary antibodies in blocking buffer. After three washes of 30 min with blocking buffer, cells were incubated with DAPI for 30 min. Finally, cells were washed for 30 min and stored in PBS. Imaging was performed briefly after the end of the immunostaining.

For immunofluorescence on spinal cords, mice were deeply anesthetized with a lethal dose of pentobarbital and perfused with 4 % PFA after blood removal with heparin. Spinal cords were collected 2 mm above and below the lesion site, post-fixed in 4 % PFA for 3 h at RT, incubated in 20 % sucrose overnight at 4°C, embedded in O.C.T. compound, and frozen at -80°C. We used 20 to 200 μm-thick cryosections. Twenty to fifty μm-thick cryosections were first submitted to antigen retrieval in citrate buffer (10 mM citrate buffer, 0.05 % Tween 20, pH 6.0) for 2 h at 65°C, washed, blocked in blocking buffer (0.1 % Triton X-100, 5% BSA, PBS) for 60 min at RT and incubated overnight at 4°C with primary antibodies diluted in blocking buffer. Sections were then washed 3 times in blocking buffer and secondary antibodies were incubated for 1 h at RT in the dark. Sections were then washed, incubated with DAPI for 10 min at RT, washed again and mounted in Citifluor (Agar Scientific). One hundred to two hundred-μm thick cryosections were permeabilized for 3 h in blocking buffer (1 % Triton X-100, 10 % FBS, PBS), and then incubated for 2 days at 4°C with primary antibodies diluted in blocking buffer. After 3 washes of 15 min with blocking buffer, sections were incubated with secondary antibodies overnight at 4°C in the dark. Sections were then washed 3 times for 15 min with blocking buffer, incubated with DAPI for 1 min and incubated overnight in 70 % Glycerol/PBS for clearing. Sections were finally washed with PBS and mounted in CitiFluor. Primary antibodies: Olig2 (Goat, 1:200, R&D Systems, AF2418), MBP (rat, 1:50. Serotec, cat. #MCA409S), CC1/APC (mouse, 1:200, Millipore, cat. #OP80), DUSP6/MKP3 (Rabbit, 1:200, Invitrogen, ARC0237, #MA5-35048), DUSP6/MKP3 (Mouse, 1:200, Santa Cruz, F-12, #sc-377070), DUSP6 (Rabbit, 1:200, Abcam, ab76310), c-Jun (rabbit, 1:200, Abcam, cat. #ab32137), c-Jun (Rabbit, 1:200, Cell signaling, 60A8, #9165), c-Jun (mouse, 1:200, BD Bioscience, cat. # 610327), Phospho-c-Jun (Rabbit, 1:100, Cell Signaling, D47G9, # 3270), Phospho-p44/42 MAPK (Rabbit, 1:200, Cell signaling, D13.1.4E, # 4370).

### Western blot analysis

Mouse sciatic nerves and primary rat OLs and primary rat SCs were lysed and processed for Western blot analysis as previously described^43^. Primary antibodies: DUSP6 (Rabbit, 1:1000, Abcam, ab76310), DUSP6/MKP3 (Rabbit, 1:350, Invitrogen, ARC0237, #MA5-35048), DUSP6/MKP3 (Mouse, 1:500, Santa Cruz, F-12, #sc-377070), c-Jun (Rabbit, 1:500, Cell signaling, 60A8, #9165), Phospho-c-Jun (Rabbit, 1:500, Cell Signaling, D47G9, # 3270), p44/42 MAPK (Mouse, 1:1000, Cell signaling, L34F12, #4696), Phospho-p44/42 MAPK (Rabbit, 1:500, Cell signaling, D13.1.4E, # 4370), Phospho-JNK1/2 (Rabbit, 1:500, Invitrogen, # 44-682G), MBP (rat, 1:750. Serotec, cat. #MCA409S), GAPDH (glyceraldehyde-3-phosphate-dehydrogenase, mouse, 1:5000, Genetex, cat. # GTX28245, lot # 821705388).

## Data Availability

The data that support the findings of this study are available from the corresponding author upon reasonable request.

## Supplementary legends

**Supplementary Figure 1. Dusp6 and phospho-ERK1/2 levels are regulated in Schwann cells and oligodendrocytes after injury. a**, Dusp6 and GFP (reporter of recombined mature OLs) co-immunofluorescence and DAPI (nuclei) labeling at 1, 5 and 7 day post spinal cord lesion (dpl) and in unlesioned (No lesion) mouse spinal cords. Representative images of 3 animals per time point are shown. Arrows point to recombined mature OLs (GFP-positive cells). **b**,**c**, Western blots of phospho-ERK1/2 (pERK1/2) and total ERK1/2 (**b**) or of Dusp6 (**c**) and quantification normalized to GAPDH at 1-3-5-12 or 2-3-4-5 days post sciatic nerve crush lesion (dpl) in crushed (Cr) compared to contralateral (Co) sciatic nerves of adult mice. Paired one-tailed (grey asterisk) or two-tailed (black asterisks) Student’s t-tests, p value: *<0.05, ***<0.001, values=mean, error bars=s.e.m., n=3 animals per time point.

**Supplementary Figure 2. Validation of Dusp6 shRNA and overexpression lentiviruses. a**,**b**, Western blots of Dusp6 and GAPDH in lysates of primary OLs transduced with lentiviruses expressing either a Dusp6-specific shRNA or a non-targeting control (Ctrl) shRNA or with control empty (Ctrl) lentiviruses or lentiviruses expressing Dusp6 (**b**). Representative images of 3 independent experiments are shown.

**Supplementary Figure 3. Dusp6 overexpression in Schwann cells leads to decreased c-Jun expression after axonal lesion**. C-Jun immunofluorescence and DAPI labeling in chamber #2 (containing SCs) of neuron/SC cultures in microfluidic devices at 1 day post axonal lesion (1 dpl) in chambers where SCs were transduced with lentiviruses expressing Dusp6 or with empty control lentiviruses. The graph shows the percentage of c-Jun-positive cells, n=3 chambers per group, 218 to 450 cells counted per chamber, unpaired two-tailed Student’s t-test, p value = *<0.05, values=mean, error bars=s.e.m.

**Supplementary Video 1. Filopodia formation and oligodendrocyte contraction around distal cut axons until their disintegration**. Live imaging (confocal imaging) of DsRed-labeled axons and Lifeact-GFP-labeled OLs in chamber #2. Time-lapse 3D surface rendering every 20 min following lesion and lasting for 24 h.

## Acknowledgements

PLP-CreERT2 mice have been used in collaboration with Dr. Ueli Suter (ETH Zürich, Switzerland). Funding: grant No. P 174 from the International Foundation for Research in Paraplegia, grants No. PP00P3_139163 and 31003A_173072 from the Swiss National Science Foundation (SNSF), grant No. JA 3019/7-1 from the Deutsche Forschungsgemeinschaft.

## Author contributions

GN and CJ conceived and designed the experiments. GN, AV, NH, KS, SLCD, JM, CB, MT and SRL performed the experiments. GN, AV, MT and CJ analyzed the data. BL, ASA, KH, DV, SB, SRL, NLJ and SMK contributed reagents/materials, GN and CJ wrote the manuscript. All authors accepted the upload of the manuscript to bioRxiv.

## Supplementary information

Supplementary Information is available for this paper.

