## Supplementary figures and images for "Repair oligodendrocytes demyelinating and disintegrating damaged axons after injury"

### Supplementary Figures

Supplementary Figure 1

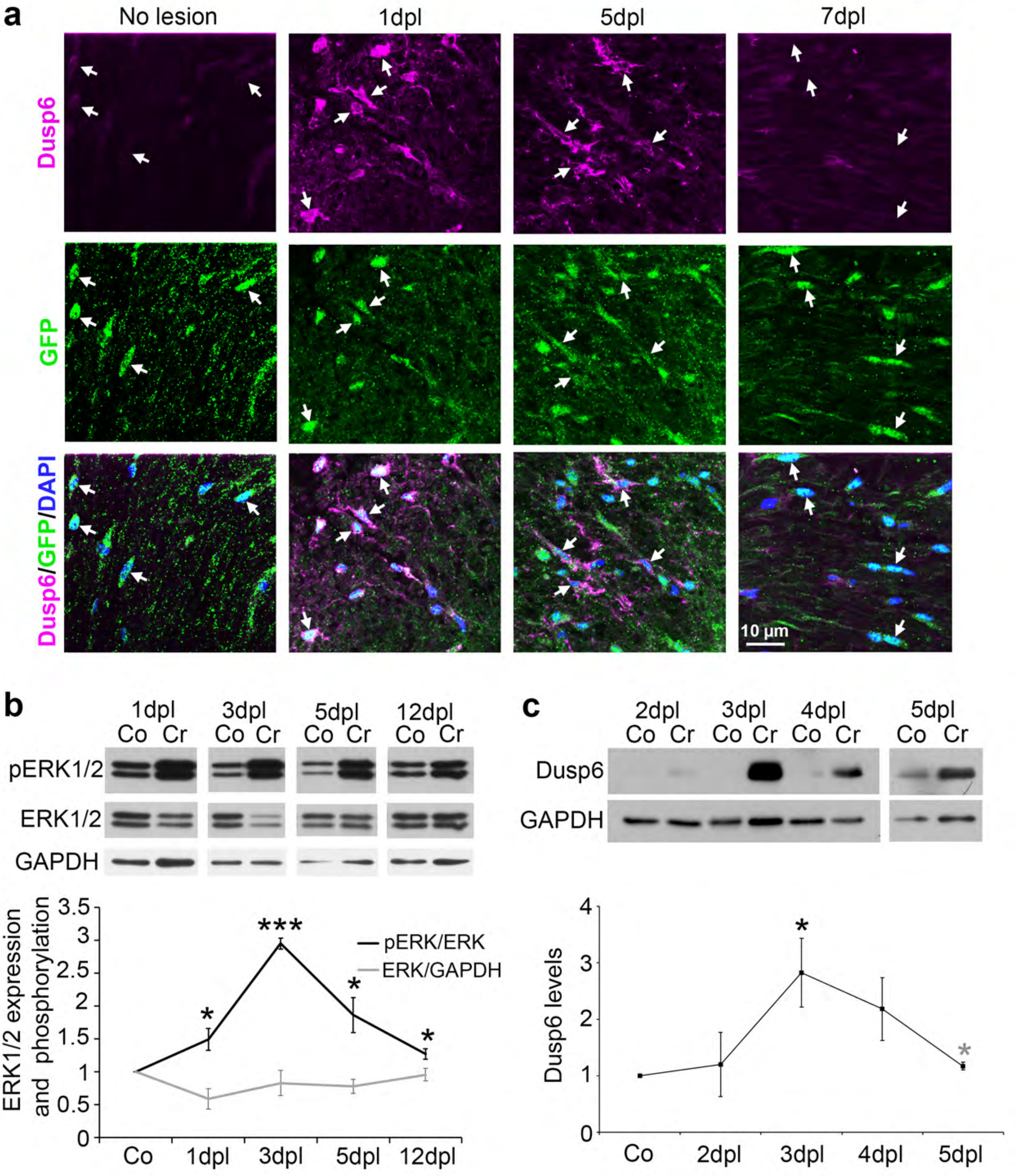

Supplementary Figure 2

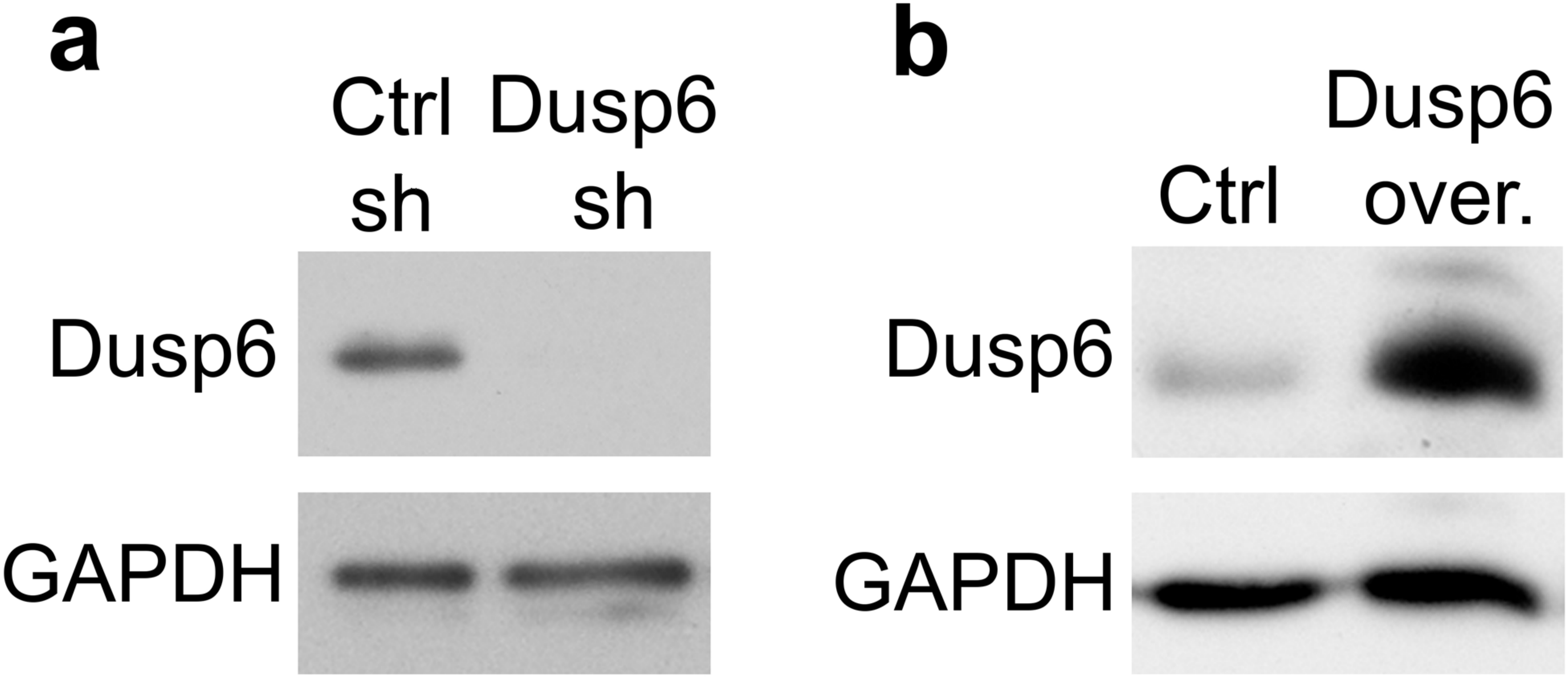

Supplementary Fig. 3

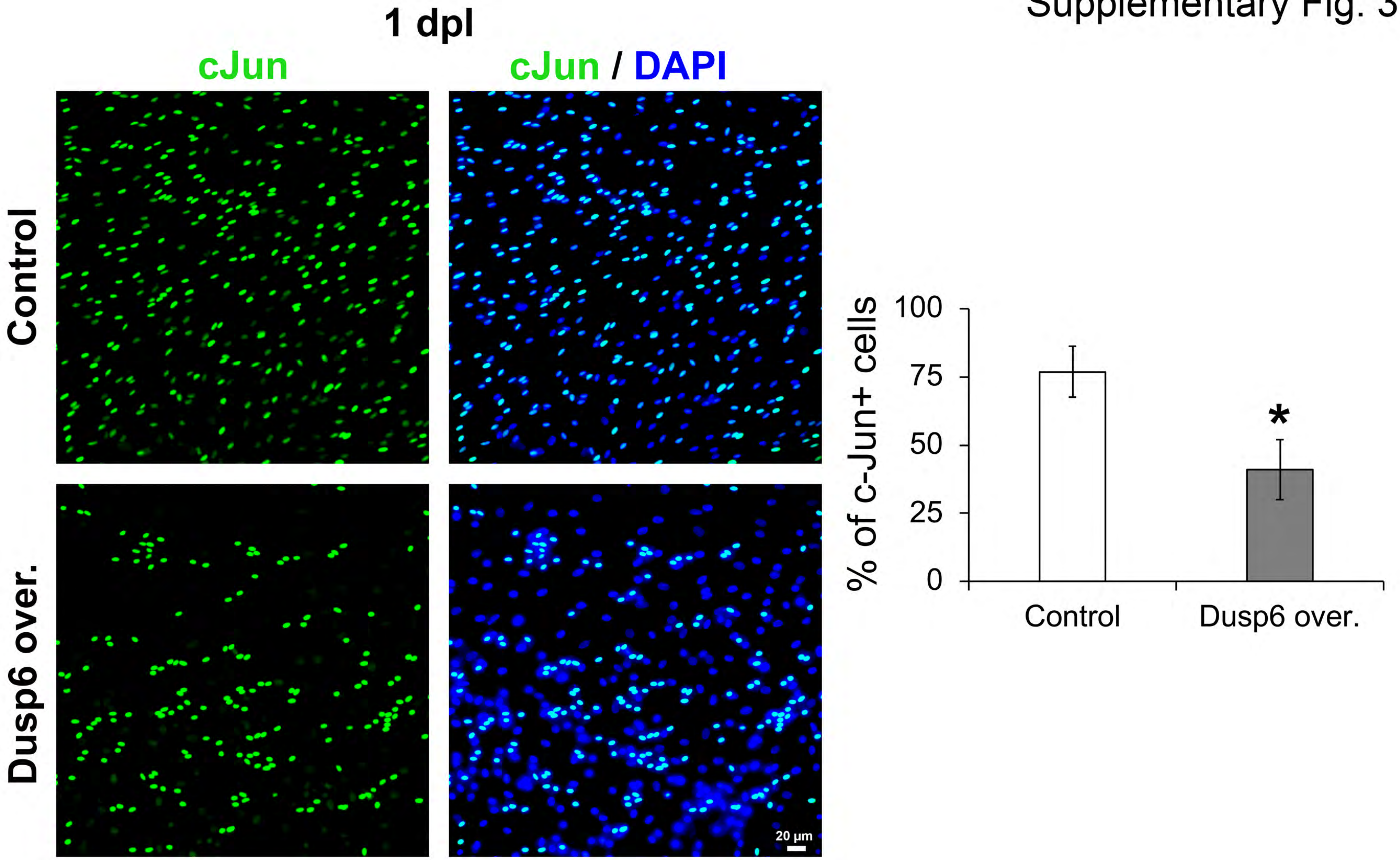
